# A transplantation experiment yields no evidence for phenotypic plasticity in shell band width in *Cepaea nemoralis*

**DOI:** 10.1101/2020.06.08.140137

**Authors:** Menno Schilthuizen, Ruth K. Scheid, Lia J. J. den Daas

## Abstract

The common European grove snail, *Cepaea nemoralis* (Helicidae), has been a model species in genetics and evolutionary biology for over a century thanks largely to its genetic shell colour polymorphism. Although most aspects of its shell colour variation are known to be purely genetic, with little or no phenotypic plasticity involved, the width of the spiral bands has been suspected to display a certain amount of plasticity. To test this, we conducted a transplantation experiment, in which 548 growing banded snails were marked and transplanted, either within the same habitat (open or closed vegetation, displaying 19% and 61% band fusion, respectively) or between habitat types. The numbers recaptured were low: 8%, 5% after removal of individuals that had not grown. Based on these samples, we did not find any substantial influence of transplantation on band width.

## Introduction

The European grove snail, *Cepaea nemoralis* (Gastropoda: Stylommatophora: Helicidae) is a well-known model species in classical genetics, ecological genetics, and evolutionary ecology (Jones et al., 1977). Its shell colour polymorphism has been used in early studies into mendelism (Lang, 1904), and later, during the wave of ecological genetics in the 1950s and 1960s, in understanding the ecological factors in the maintenance of genetic polymorphism and microevolution (e.g., Cain & Sheppard, 1950, 1954; Lamotte, 1959; Wolda, 1969a). More recently, its popularity has been revived because of its use in evolutionary citizen science projects (Silvertown et al., 2011; Worthington et al., 2012; Kerstes et al., 2019) and because its value in measuring the evolutionary impact of climate change was recognised (Ożgo & Schilthuizen, 2013, Ożgo et al., 2017).

Although the value of this species as a model is helped by its size, conspicuous colours, abundance, wide distribution and wide ecological tolerances, not least among its redeeming features is the fact that its shell colour polymorphism appears to be fully genetic, with little or no influence of phenotypic plasticity (Lang, 1904; Cook, 1967; Wolda, 1969b; Murray, 1975; Kerkvliet et al., 2017; Davison et al., 2019).

The following is based on Murray (1975). In *C. nemoralis*, the locus *C* is responsible for the ground colour of the shell, and different alleles determine colours that range from pale yellow to deep brown. In addition, the shell can display between one and five dark brown bands, which may merge with one another. The presence or absence of bands is controlled by the locus *B*. The dominant allele codes for absence of bands, the recessive allele for their presence. At the *T* locus, the dominant allele removes the top-most two bands, leaving only the three lowest. The dominant allele of the *U* locus suppresses bands 1, 2, 4, and 5, leaving a mid-banded shell. Five other loci (I, S, P, R, and O) affect various aspects of band pigmentation, width, and evenness. Phenotypes are denoted with a letter (Y, P, or B) for the shell ground colour (yellow, pink, and brown), followed by five digits (1-5), each replaced by a zero when a band is lacking. Band fusion is coded with parentheses around the conjoined band digits. A yellow, banded shell with bands 1, 2, and 3 fused, and band 4 missing, for example, is coded as Y(123)05.

Although full genetic control of single loci has been demonstrated for most aspects of shell colour and banding, one feature, namely band width, has remained somewhat elusive. It is suspected to be under polygenic control (Murray, 1975), and may also be affected by a degree of phenotypic plasticity (R. Cameron, personal communication; Schilthuizen & Kellermann, 2013). In this paper, we conduct transplantation experiments in *C. nemoralis* to investigate whether any indication for phenotypic plasticity in band width can be demonstrated.

## Materials and methods

The experiment was carried out near the ‘Kathedralenbos’ (52.323°N 5.318°E), a land-art project near Almere, Prov. Flevoland, the Netherlands. Previously, Schilthuizen (2013) showed that *Cepaea nemoralis* at this locality differ strongly in band width, depending on the habitat. In shaded habitats, 61% of individuals had fused bands, whereas in open habitats, only 19% of the population did. In four sites (named Shaded-1 and Shaded-2, and Open-1 and Open-2; see Fig. 1), in June and August, 2013, we collected living, juvenile, banded snails of intermediate size (10-15 mm diameter). Shaded sites consisted of closed-canopy deciduous forest, whereas open sites were tall grasses and herbs in a road-side verge.

**Fig. 1.**
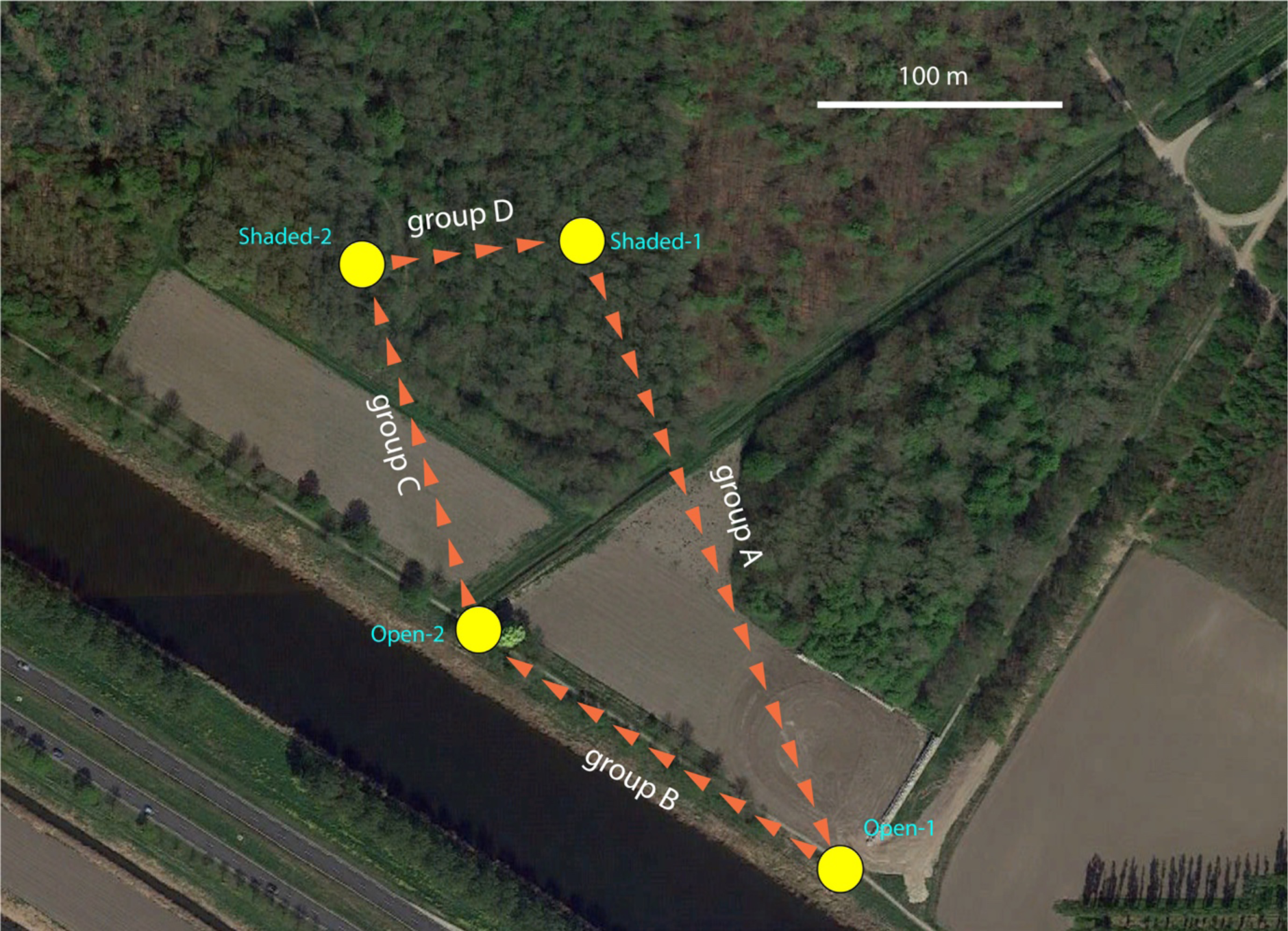
The vicinity of the Kathedralenbos, with the four sample sites and the transplantations indicated.

Sample dates and sizes were as follows. Shaded-1 [A]: June 17^th^ (N=56), August 13^th^ (N=44); Open-1 [B]: June 17^th^-18^th^ (N=60), August 13^th^ (N=85); Open-2 [C]: June 19^th^ (N=60), August 13^th^ (N=118); Shaded-2 [D]: June 18th (N=61), August 13^th^-14^th^ (N=65). Total sample sizes were therefore 100, 145, 178, and 126 for Shaded-1 [A], Open-1 [B], Open-2 [C], and Shaded-2 [D], respectively.

Each snail was marked (using black Indian ink, either directly onto the shell or on a dot of white correction fluid that had been applied first) with an individual code consisting of a capital A, B, C, or D (representing the source site / transplant group), and a unique number. In addition, a dot or vertical line was drawn on the shell, 4 mm in apical direction from the edge of the aperture (Fig. 2). The markings were covered with a layer of clear nail varnish. Each snail thus marked was photographed, measured, and then released. Releases were carried out as indicated in Fig. 1, which ensured that two transplantations were within the same habitat type (Shaded-2 to Shaded-1 and Open-1 to Open-2), and the other two transplantations were into the alternative habitat (Shaded-1 into Open-1 and Open-2 into Shaded-2). Releases were carried out within a circle with a radius of 20 m, in the centre of the area where the snails from that site were collected.

**Fig. 2.**
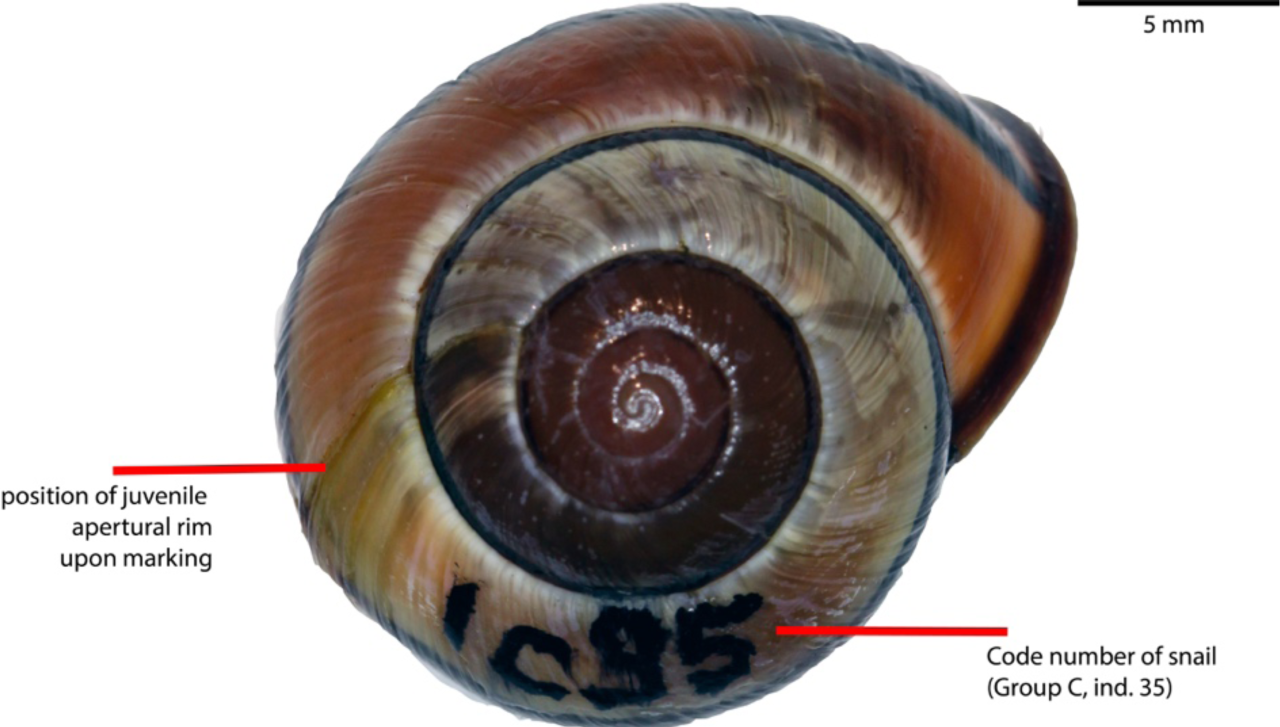
An adult snail from group 3, having been marked as a juvenile, and showing the code number and the stripe-mark 4 mm downstream from which the juvenile apertural margin had been located.

For recapturing, the four sites were visited three times: on August 17th and 18th, 2013 (during which snails released in June, 2013, were recaptured), on June 26th, 2014 (during which snails released in June and August, 2013, were recaptured), and on August 8th, 2014 (during which snails released in June and August, 2013, and June, 2014, were recaptured). Of all retrieved snails (whether they were dead or alive), we measured the band widths at the position where the black line had been drawn before release, and also all along the whorl that had accrued after release, from 0 to 10 mm, with 2 mm intervals.

## Results and Discussion

Overall, we retrieved 3, 11, 15, and 15 marked snails at sites Shaded-1, Open-1, Open-2, and Shaded-2, respectively (Supplementary Table 1), an overall recapture rate of 8%. However, a portion of these had not grown at all or very little (< 2 mm), either because they had presumably died shortly after marking, or because they had been marked at the developmental stage just prior to reaching maturity. After removing those individuals, we were left with 1, 9, 12, and 6 individuals from sites Shaded-1, Open-1, Open-2, and Shaded- 2, respectively, that could be used to assess phenotypic plasticity.

Our measurements (Supplementary Table 1, Fig. 3) show that band widths did not change appreciably, neither in the experimental groups (transplanted to a different habitat type) nor in the control groups (transplantation within the same habitat type). In most individuals, a growth halt was visible at the moment the animal had been marked, probably as a result of the shock of capture, handling, and release in a new environment, and bands tended to be more weakly pigmented immediately after growth recommenced. However, this loss of pigmentation (which is commonly seen after growth halts in *Cepaea*) was usually of short duration, and in most cases the original band width and darkness were reinstated quickly. We did not find any individuals in which band width changed abruptly and permanently after transplantation.

**Fig. 3.**
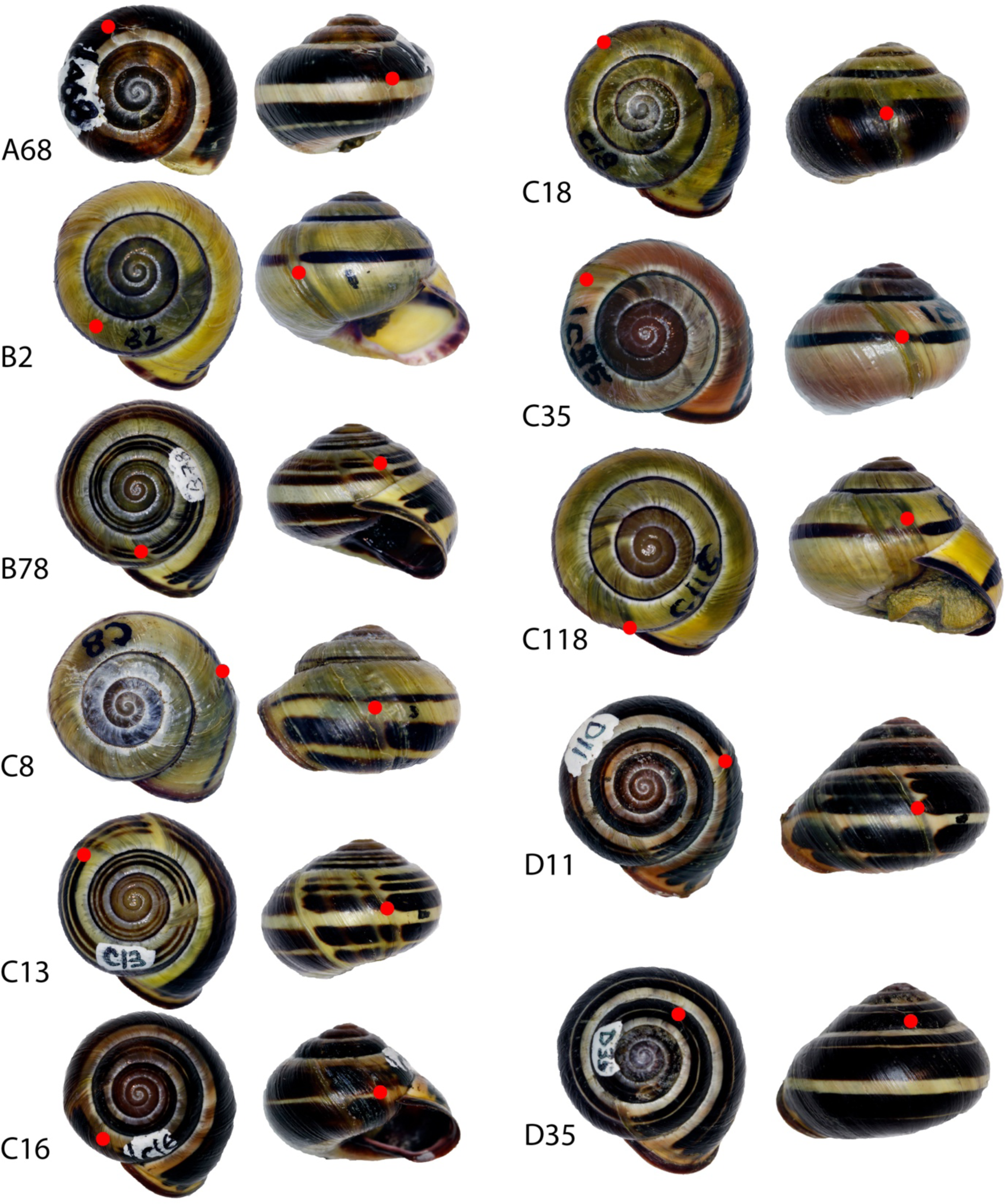
A representative set of marked and recaptured snails that had grown a substantial amount (dorsal and lateral views), individual codes given to the left of each dorsal view. The apertural rim upon marking is indicated with a red dot; everything downstream from this red dot has grown after transplantation. Groups A and C have been transplanted across habitat types, Groups B and D within habitat types.

Our results suggest that, like all other shell colour traits in *Cepaea*, band width is purely genetic and not or not appreciably under environmental control. We should note, though, that our samples of retrieved snails are small, and in one experimental group (group A, which was transplanted from Shaded to Open habitat), we retrieved only a single snail. Also, our experiment was carried out in a single population. To confirm these results, the experiment should be replicated in multiple locations, and preferably even larger sample sizes, given the relatively low recapture rates in our study.

## Acknowledgements

For assistance during the field work, we thank Monica Lodi and Heike Kappes.

